# JUUL and Combusted Cigarettes Comparably Impair Endothelial Function

**DOI:** 10.1101/754069

**Authors:** Poonam Rao, Jiangtao Liu, Matthew L. Springer

## Abstract

**Objectives:** JUUL and earlier generation electronic cigarettes (e-cigs) are promoted as being less hazardous than cigarettes. While JUUL Labs, in particular, claims that switching from smoking to vaping has beneficial impacts, the health effects of such products are not well understood. We investigated whether exposure to JUUL and previous generation e-cig aerosol impairs endothelial function comparably to cigarette smoke.

**Methods:** We exposed rats to aerosol from Virginia Tobacco flavor JUUL, an e-cig tank system using unflavored freebase nicotine e-liquid, Marlboro Red combustible tobacco cigarettes, or clean air for 10 cycles of 2 second inhalation over 5 minutes. Endothelial function (FMD) was measured pre- and post-exposure. Blood was collected 20 mins post-exposure for serum nicotine analysis.

**Results:** Aerosol/smoke from JUUL, previous generation e-cigs, and cigarettes all impaired FMD. The extent of impairment ranged from 34%-58%, although the differences between groups were insignificant. Nicotine was highest in serum from the JUUL group; for the other e-cig and cigarette groups, nicotine levels were lower and comparable to each other.

**Conclusions:** Aerosol from JUUL and previous generation e-cigs impairs endothelial function in rats, comparable to impairment by cigarette smoke.

## INTRODUCTION

It has only been a few years since e-cigarettes (e-cigs) became a major player on the tobacco product landscape,^1^ but the disruption in the tobacco world has been undeniable. If the original appearance of e-cigs was a sea change in how tobacco products are used, the field has now experienced a seismic event with the arrival of JUUL and similar pod-based products using nicotine salts being the most recent product innovations. Since 2016, there has been a dramatic increase in youth e-cig use,^2^ with JUUL devices particularly effective at recruiting teenagers to start nicotine usage.^3^

JUUL is new, and its effects on health relative to combusted cigarettes and earlier versions of e-cigs are unclear. As with earlier generation e-cigs, the e-liquid in JUUL pods is composed of vegetable glycerin (VG) and propylene glycol (PG) along with flavors and nicotine. However, while the freebase nicotine used in earlier generations of e-cigs limits the amount that can be comfortably inhaled, JUUL introduced the use of acidified nicotine salts that are tolerated by the airway epithelium, leading to the ability to deliver nicotine at substantially higher concentrations.^4^ There is no single defining characteristic that distinguishes between JUUL and previous generation e-cigarettes, which differ based on nicotine concentration, nicotine molecular form, and device design; therefore, we are using the term “Previous generation e-cig” to refer to free base nicotine e-liquid in a Nautilus Aspire tank. The high concentration of nicotine salts has been implicated in the youth epidemic.^5^ Much of the current literature focuses on the effect of non-aerosolized e-liquid or aerosol condensate on cultured cells,^6^ so there is a need for integrative physiological assessments of JUUL’s effects on health as the FDA grapples with what they and the Surgeon General are now describing as a youth vaping epidemic.^7,8^

Endothelial function assessed as arterial flow-mediated dilation (FMD) is a validated measure for overall cardiovascular health.^9–11^ It is impaired in humans by smoking of combusted cigarettes^12^ and chronic or acute exposure to secondhand smoke.^13–15^ Our micro-ultrasound-based approach to measure FMD in living rats^16^ yields results whose pharmacological and biophysical effects are similar to those observed in humans.^12,15^ Using this technique, we showed that FMD is also impaired in rats by brief exposures to mainstream and sidestream smoke or mainstream aerosols from cigarettes, little cigars, combustible marijuana, and IQOS “heat-not-burn” tobacco products.^17–20^ Acute use of e-cigs with or without nicotine in humans impairs several measures of endothelial function including FMD^21,22^ and we have reported that nitric oxide production from cultured endothelial cells is impaired when the cells are incubated in serum from chronic e-cig use.^23^ Here we report that acute mainstream exposure to aerosol from JUUL or from previous generation e-cigs using freebase nicotine impairs vascular function comparably to combusted cigarette smoke and delivers considerably more nicotine to the blood on a per puff basis.

## METHODS

### Animals

We used Sprague-Dawley rats, 10 weeks old, n=8 rats/group (4 male, 4 female) with body weight of 300-350 g (males) or 185-220 g (females), as has been the standard condition for our previous studies on smoke exposures.^17–20^ Rats were anesthetized with intraperitoneal injection of ketamine (100 mg/kg) and xylazine (5 mg/kg). During the experiment, rats were kept on a thermal pad to maintain temperature 37.5°C and prevent hypothermia. Frequency and depth of respiration was continuously monitored to ensure full anaesthesia and supplemental intramuscular anaesthetic was given if necessary. All procedures were approved by the UCSF Institutional Animal Care and Use Committee.

### Measurement of Endothelial Function

FMD was performed as described previously.^16^ Briefly, after anesthesia as described above, we made a 1 cm incision in the rat’s groin and surgically exposed the right common iliac artery. We then placed an arterial loop occluder consisting of a 4–0 Prolene filament under the artery and passed it through a 15 cm PE–90 tubing to enable transient occlusion of blood flow after suturing the skin. A series of diameter images of the femoral artery and accompanying Doppler blood flow images were recorded using Vevo660 ultrasound system with a 35 MHz transducer (VisualSonics, Toronto, Canada) before occlusion. Then we induced a transient occlusion for 5 minutes followed by release of the snare to re-establish perfusion with a rush of blood flow (hyperemia), with ultrasound measurements of femoral artery diameter performed every 30 seconds for 3 minutes with additional measurements at 4 and 5 minutes.

FMD was measured before and after exposure, as summarized schematically in Figure 1. We used an automated program (Brachial Analyzer 5; Medical Imaging Applications, Coralville, Iowa, USA) to measure baseline artery diameter and peak post-ischemia diameter during diastole. The investigator was blinded to exposure conditions during FMD procedure, analysis of ultrasound images, and subsequent calculations. FMD was calculated as % change: (peak diameter_postischemia_ − diameter_baseline_)/diameter_baseline_×100.

**Figure 1.**
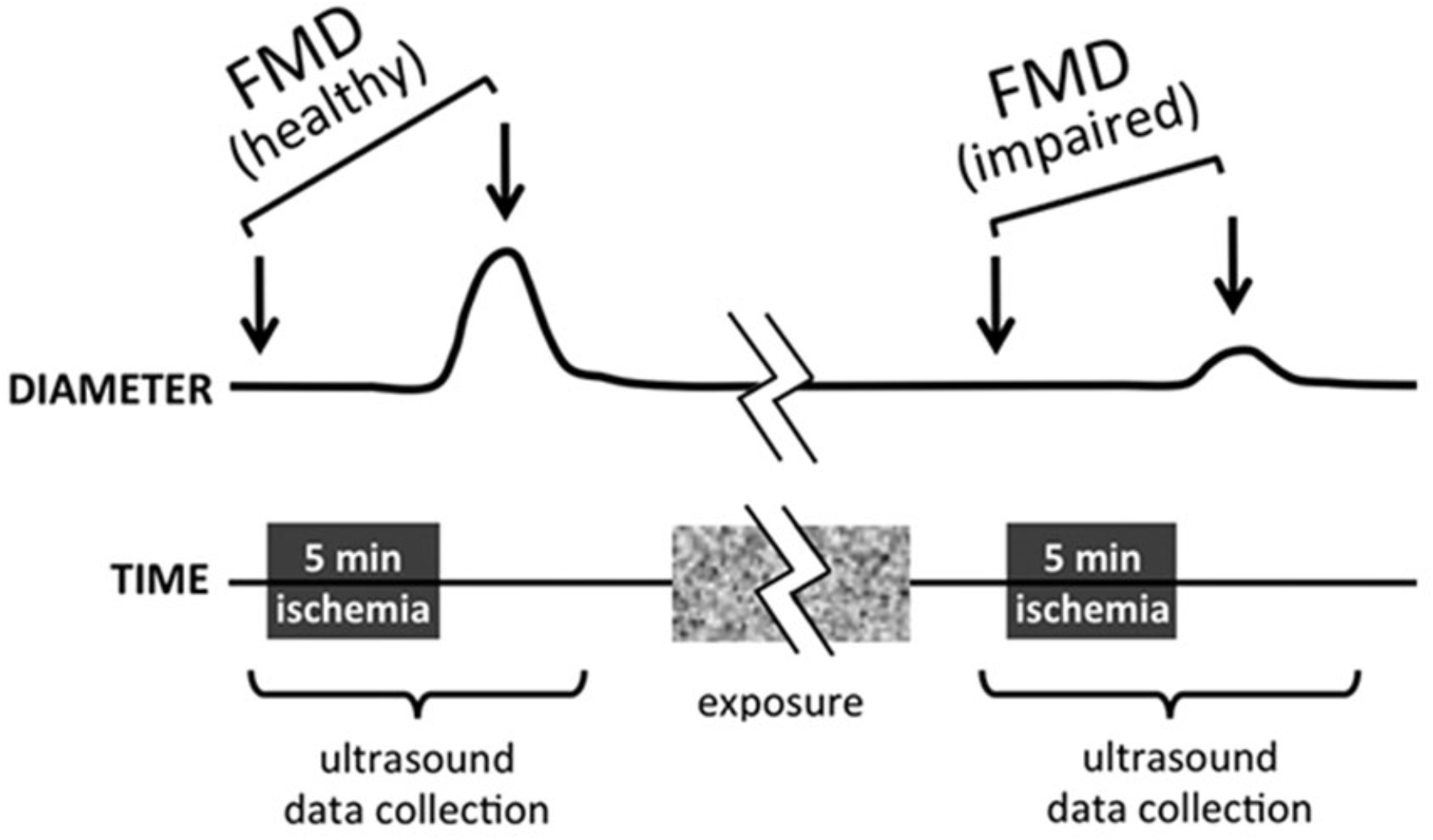
Schematic diagram showing time course of ultrasound readings during FMD measurement, pre- and post-exposure.

### Mainstream Smoke/Aerosol Generation and Exposure

A JUUL starter kit was purchased from a rural California gas station mini-mart. To represent previous generation e-cig aerosol, a Nautilus Aspire tank and e-liquid components were acquired from MyFreedomSmokes. Cigarettes were Marlboro Red brand. Both the freebase nicotine e-liquid and the cigarettes were un-flavored, and JUUL pods were Virginia Tobacco flavor. (There are no unflavored JUUL pods, but Virginia Tobacco flavor is reported to have the lowest level of flavorants of the 8 flavors on the market as of August 2019.^24^)

We exposed anesthetized rats via nose cone to aerosol from “5% nicotine” JUUL pods (listed percent nicotine is by weight but corresponds to 59 mg/ml; JUUL e-liquid is 60% VG and 30% PG by volume), Nautilus e-cig tank using freebase nicotine e-liquid (“Previous generation e-cig”; 33% VG, 67% PG, 12 mg/ml nicotine), Marlboro Red cigarettes as a positive control, and clean air as a negative control (Figure 2). The exposure regimen consisted of a series of consecutive 30s cycles, each including 2s of inhalation, over 5 min. To generate the aerosol and mainstream smoke, we used a Gram Universal Vaping Machine version 5.0 (Gram Research, Oakland, California, USA). The vaping machine contains an automatic syringe pump and has been shown to reproducibly generate aerosols from e-cigarettes under controlled conditions.^25^ The system generated 35 ml of aerosol over 2 s; thus, the cigarette smoke generation was consistent with the ISO standard 3308:2012 smoking regimen for each puff but the frequency was doubled to remain directly comparable to the e-cig exposure timing.

**Figure 2.**
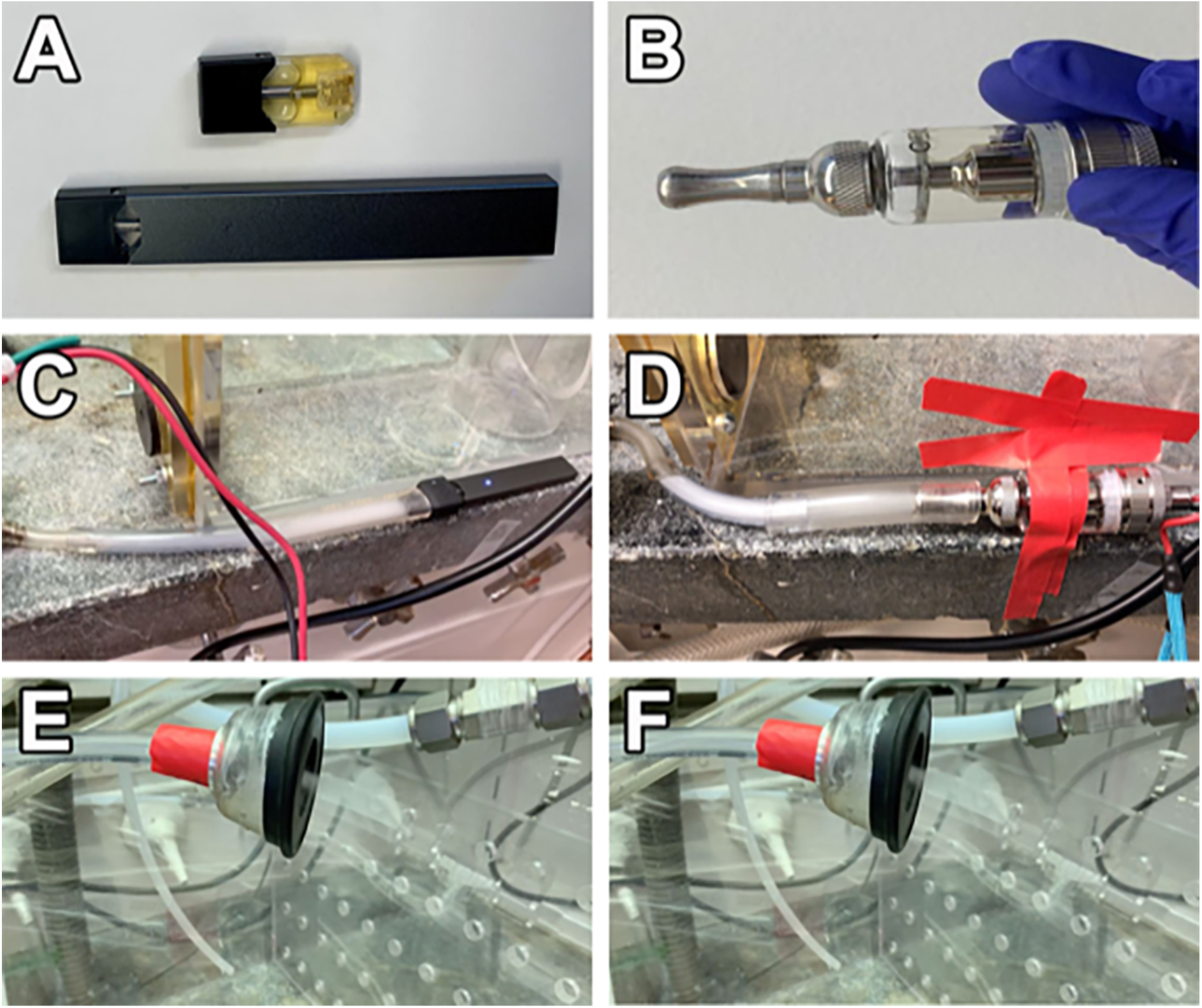
JUUL and previous generation e-cig used for this study. A) JUUL stick with loaded pod on the bottom, Virginia tobacco JUUL pod on top. B) Nautilus Aspire tank with freebase nicotine e-liquid. C) JUUL attached to Gram system. D) Nautilus Aspire tank attached to Gram system. E) JUUL aerosol from nose cone. F) Freebase nicotine e-cig aerosol from nose cone.

Particle levels were not controlled because we used whatever mainstream aerosol or smoke was generated by a simulated inhalation through these products. Separate syringes, valves, and tubing segments were used to avoid cross-contamination of different aerosols and smoke.

Rats from each group were exposed in a random order and arterial diameter measurements were obtained by an investigator unaware of the experiment condition.

### Statistics

To evaluate differences in FMD or baseline diameter versus time or exposure conditions, we fit a two-factor (exposure condition and time) repeated measures ANOVA to all data using a linear mixed model estimated with restricted maximum likelihood estimation, then tested for differences over time and across exposure conditions using contrasts and pairwise comparisons, adjusted for multiple comparisons using the Šidák method using STATA 13.1.

## RESULTS

### Exposure to Aerosol from JUUL and Previous Generation E-cigs Impaired FMD Comparably to Cigarette Smoke

We initially conducted an n=4, single group pilot experiment to test our adaptation of the vaping machine to JUUL and to confirm that our exposure regimen of 10 puffs over 5 minutes was sufficient to observe impairment of FMD. FMD declined significantly from 6.4±1.5% (SD) to 3.2±1%, p=.04 by 2-tailed paired t-test even with this small group size.

Using the same conditions with groups of n=8 in a blinded experiment comparing our four main exposure conditions, FMD was impaired by exposure to JUUL aerosol (8.6±1.5% pre-exposure vs. 3.6±0.8% post-exposure, p=.003), previous generation e-cig aerosol (8.8±0.8% pre-exposure vs. 5.3±0.6% post-exposure, p=.001), and Marlboro Red cigarette smoke (8.8±1.4% pre-exposure vs. 5.8±1.0% post-exposure, p=.03) (Figure 3). The extent of FMD impairment did not significantly differ between these groups (p=0.83). No significant impairment of FMD was seen in the air group (7.1±0.5% vs. 6.4±1.1%, p=.54).

**Figure 3.**
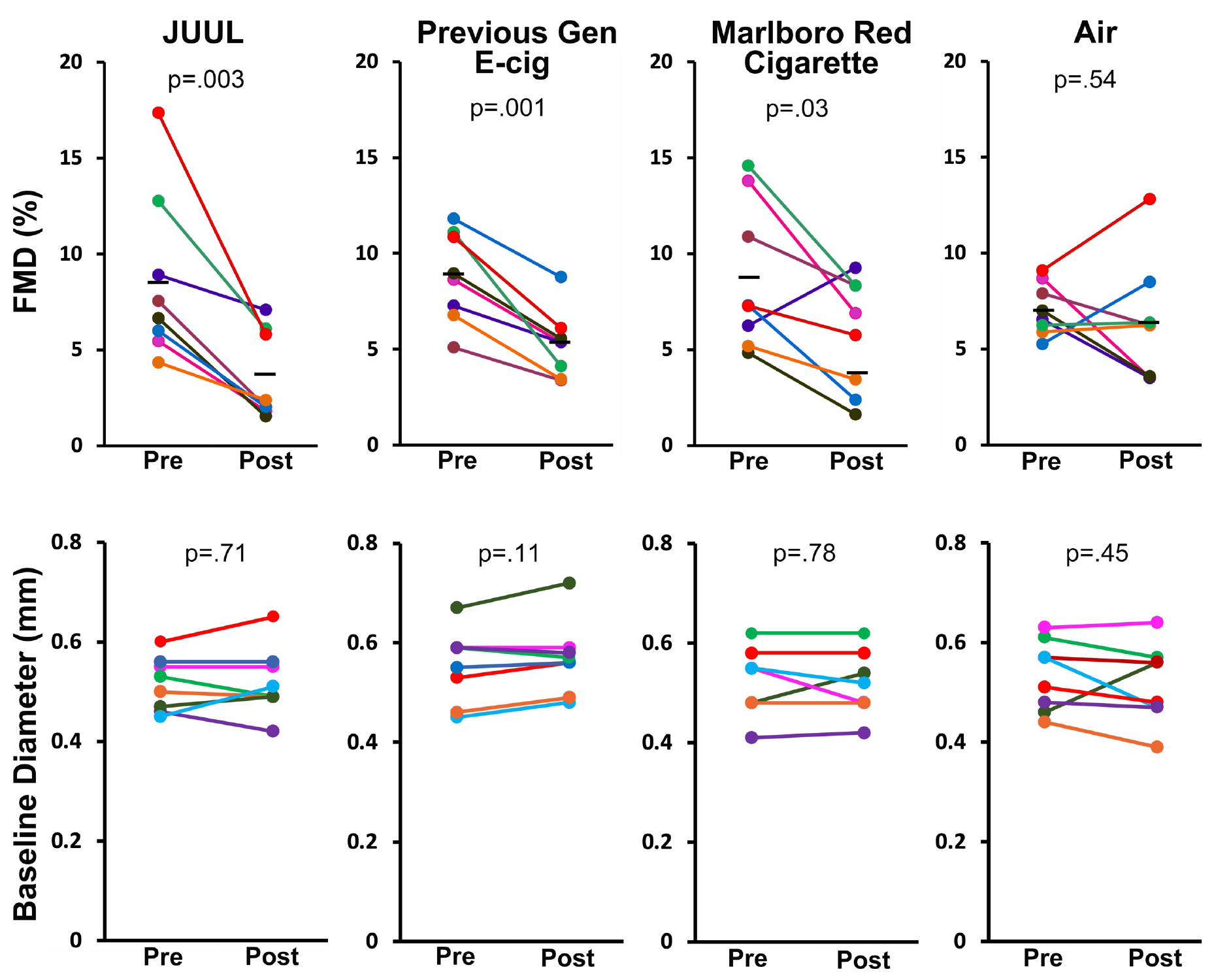
FMD was impaired by JUUL aerosol, previous generation e-cig aerosol, and Marlboro Red cigarette smoke. Top: FMD after 5 mins of exposure. Bottom: Baseline arterial diameter (before arterial occlusion). Colored lines denote individual rats. Horizontal black bars denote the mean of the respective groups. p values are derived from paired 2-tailed t tests. “Previous Gen” = previous generation.

### Serum Nicotine was Highest in Rats that were Exposed to JUUL Aerosol

In rats exposed to JUUL aerosol, serum nicotine level from samples collected 20 mins after end of exposure was 136.4±23.6 ng/ml, which was significantly higher (p<0.001) than mean nicotine levels in the previous generation e-cig group (17.1±4.9 ng/ml) and in the cigarette group (26.1±4.2 ng/ml) (Figure 4). Similarly, serum cotinine levels in the JUUL group (41.5±8.7 ng/ml) were significantly higher than the previous generation e-cig (8.3±2.5 ng/ml) and cigarette groups (14.1±1.0 ng/ml). Serum nicotine and cotinine in the air group were undetectable.

**Figure 4.**
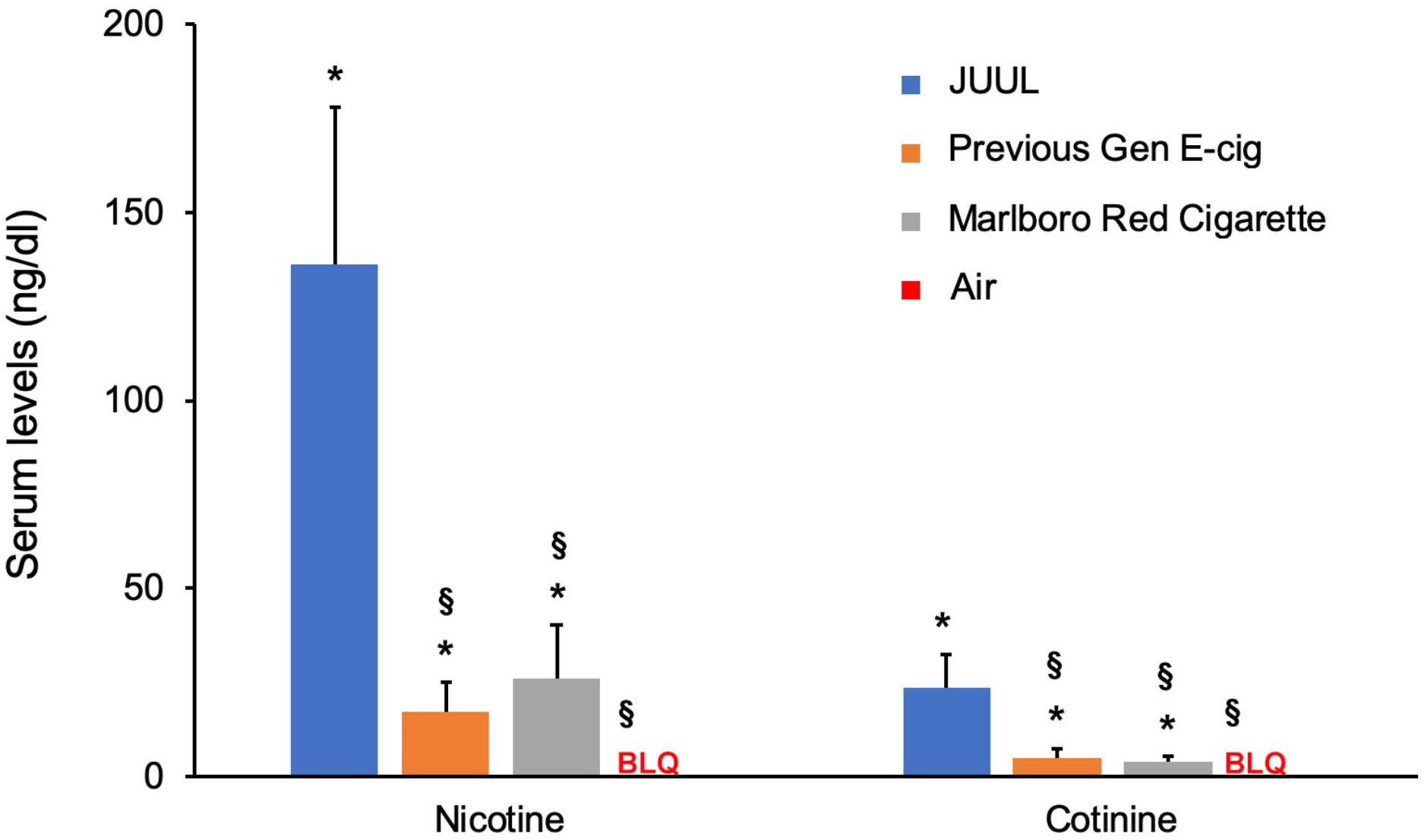
Serum levels (ng/ml) of nicotine and cotinine from sera collected after 20 mins of exposure. *p<0.001 compared to air group. §p<.001 compared to JUUL. BLQ = below limits of quantitation. “Previous Gen” = previous generation.

## CONCLUSIONS

Carnevale et al.^21^ previously reported that exposure to previous generation e-cig aerosol leads to adverse effects on oxidative stress and vascular function in humans, and reports of harmful effects on hemodynamics and arterial stiffness are accumulating.^22,26,27^ Our study focuses on the effect of acute exposure to JUUL on vascular function. Our results in this rat model indicate that brief use of a JUUL, a previous generation e-cig, or a Marlboro Red cigarette all led to comparable impairment of endothelial function, suggesting that the adverse effect of cigarettes on vascular endothelial function that has been a known consequence of cigarette smoking since the 1990s^12,15^ is not prevented by use of JUUL.

Despite comparable drops in FMD after 10 puffs over 5 minutes, the rats exposed to JUUL aerosol absorbed a much larger dose of nicotine than those exposed to the same number of puffs of previous generation e-cigarette aerosol or of cigarette smoke. This is consistent with the fact that JUUL e-liquid delivers a much higher concentration of nicotine salts than that of freebase nicotine found in previous generations of e-cigs, which enables the device to use lower battery power than tank models and vape pens that use freebase nicotine. A potential advantage of this approach is that the lower power is expected to generate fewer chemical breakdown products or liberated metals at the heating coil, and lower levels of particulate aerosols. However, results from our study show that this property of JUUL did not prevent the substantial reduction of endothelial function.

It is worth considering whether the higher amount of nicotine absorption by the rats per inhalation from JUUL is reflective of what occurs in human users. This may depend on the population in question. The unconscious rats inhaled 10 cycles of aerosol or smoke as they were administered, but humans control their self-administration of nicotine based on feedback as it is absorbed. Adult former cigarette smokers who are conditioned to a specific level of nicotine may stop their JUUL vaping session when that state has been achieved. However, adolescent non-smokers who are not familiar with the effects of nicotine may be more likely to chase higher levels of the drug’s effects, and the ease of over-consuming nicotine with JUUL makes this likely, especially in light of reports of teenagers binging on JUUL to the point of rapid addiction and behavioral consequences.^3,28,29^

It is likely, however, that the difference in nicotine absorption from JUUL and conventional e-cig aerosol does not directly dictate the magnitude of the vascular response, which we have already observed to be saturating in this model even by very brief (2 seconds) mainstream exposures to cigarette smoke and IQOS aerosol.^20^ While we have reported a positive correlation between the concentration of nicotine in heavily diluted tobacco sidestream smoke and the extent to which FMD is impaired in rats exposed to it,^30^ the fact that FMD is still impaired in humans inhaling nicotine-free e-cig aerosol^22^ and in rats inhaling marijuana sidestream smoke^19^ indicates that the nicotine is not the main driver of this effect.

A limitation of this study is that it cannot directly determine on a per unit basis whether aerosol from JUUL or freebase nicotine e-cigs is less inhibitory to vascular function than cigarette smoke. We did not do a dose-response comparison, because smokers and vapers inhale whatever is delivered from the product being used, without dilution, making the question of relative inhibitory effects academic. It is likely that people who inhale e-cig aerosol regardless of the source, or cigarette smoke from active smoking, are saturating the effect on endothelial function and thereby receiving equivalent magnitudes of physiological consequences.

## IMPLICATIONS FOR TOBACCO REGULATION

The comparison of cardiovascular health effects of JUUL use with those of previous generation e-cigs and of combusted cigarettes is an important issue for policymakers, including the FDA and comparable bodies outside the US. This a timely issue, as the FDA is currently grappling with the explosion of youth use of JUUL amid the frequent assumption among its users that JUUL is relatively harmless. A larger study of the relative contributions of JUUL e-liquid components to endothelial dysfunction (e.g., PG and VG concentrations, nicotine state of freebase vs. salts, popular JUUL flavors like Mango) is underway, but our initial main finding is of timely importance, so we are publishing this Brief Report to make the results available now. Our finding about comparable endothelial toxicity of aerosols from JUUL and previous generation e-cigs, and combusted cigarette smoke, indicates that, at least for this important cardiovascular outcome, modified risk claims are not appropriate for e-cigarettes, including JUUL.

## Acknowledgements

We thank Drs. Stanton Glantz, Bonnie Halpern-Felsher, Leila Mohammadi, and Xiaoyin Wang for helpful comments on this manuscript. Research reported in this publication was supported by grant R01HL120062 from the National Heart, Lung, and Blood Institute at the National Institutes of Health (NIH/NHLBI) and the US Food and Drug Administration Center for Tobacco Products (FDA CTP); grant U54 HL147127 from the NIH/NHLBI and FDA CTP; and a generous donation from the Elfenworks Foundation in memory of Deb O’Keefe. The content is solely the responsibility of the authors and does not necessarily represent the official views of the NIH or the FDA. This manuscript has been posted on the bioRxiv preprint server.

